# Brain networks are decoupled from external stimuli during internal cognition

**DOI:** 10.1101/2021.01.18.427211

**Authors:** Dror Cohen, Tomoya Nakai, Shinji Nishimoto

## Abstract

Our cognition can be directed to external stimuli or to internal information. While there are many different forms of internal cognition (mind-wandering, recall, imagery etc.), their essential feature is independence from the immediate sensory input. This is thought to be reflected in the decoupling of brain networks from the external stimuli, but a quantitative investigation of this remains outstanding. Here we present a conceptual and analysis framework that links stimulus responses to connectivity between brain networks. This allows us to quantify the coupling of brain networks to the external stimuli. We tested this framework by presenting subjects with an audiovisual stimulus and instructing them to either attend to the stimulus (external task) or engage in mental imagery, recall or arithmetic (internal tasks) while measuring the evoked brain activity using functional MRI. We found that stimulus responses were generally attenuated for the internal tasks, though they increased in a subset of tasks and brain networks. However, using our new measure of coupling, we showed that brain networks became increasingly decoupled from the stimulus, even in the subset of tasks and brain networks in which stimulus responses increased. These results quantitatively demonstrate that during internal cognition brain networks are decoupled from external stimuli, opening the door for a fundamental and quantitative understanding of internal cognition.

## Introduction

A fundamental property of cognition is that it can be directed either to external stimuli (external cognition) or to internally generated information (internal cognition). For example, during a commute to work our thoughts may be directed at the landscape passing us by or to pleasantly imagining our next holiday. Though the majority of studies to date have focused on external cognition, it is estimated that as many as 50% of our waking thoughts are internally directed (Schooler et al., 2011). The remarkable prevalence of such thoughts, combined with potential connections to clinical conditions such as depression and anxiety, has led to increased focus on internal cognition (Andrews-Hanna et al., 2014; Christoff et al., 2016; Perkins et al., 2015; Zabelina & Andrews-Hanna, 2016).

One difficulty with studying internal cognition is that it is a broad phenomenon, characterized along several dimensions. For example, internal cognition can involve different sensory modalities (e.g. aural vs visual imagery) and temporal orientations (e.g. recall vs prospective planning), and can be either spontaneous (e.g. mind-wandering, day-dreaming) or intentional (e.g. cued imagery or recall)(Andrews-Hanna et al., 2014; Christoff et al., 2016; Dixon et al., 2014; Smallwood, 2013; Smallwood & Schooler, 2015). However, a common feature of all these forms of internal cognition is that they require a shift from the immediate external stimuli to internally generated information, referred to as perceptual decoupling (Smallwood & Schooler, 2015).

Perceptual decoupling is thought to decouple the brain process that support internal cognition from the external stimuli (Christoff et al., 2016). To this end, a number of electroencephalogram (EEG) studies have demonstrated attenuated stimulus responses (amplitude or phase locking) during episodes of mind wandering (Baird et al., 2014; Barron et al., 2011; Kam et al., 2011; Schooler et al., 2011; Smallwood et al., 2008), as well as reductions in the reliability of neural responses during an internal task (mental arithmetic) (Ki et al., 2016). However, these studies did not link the reduction in responses to the brain processes that support internal cognition. Thus, a quantitative demonstration that brain processes are decoupled from external stimuli during internal cognition remains outstanding.

To investigate the decoupling of brain processes from external stimuli we first need to identify the brain process that support internal cognition. There is a growing literature emphasizing that internal cognition is supported by specific connectivity patterns between brain areas. In particular, interactions within and between the default mode (DMN), Attention and Executive networks are considered fundamental to internal cognition (Andrews-Hanna et al., 2014; Benedek et al., 2016; Buckner & DiNicola, 2019; Dixon et al., 2014; Kam et al., 2019; Smallwood et al., 2012; Turnbull et al., 2019; Zabelina & Andrews-Hanna, 2016). In a more concrete example, Shirer and colleagues showed that functional magnetic resonance imaging (fMRI) connectivity patterns very reliably distinguish between different forms of internal cognition (mental arithmetic, episodic memory and recall (Shirer et al., 2012)). Thus, to test whether brain processes are decoupled from the external stimulus it is necessary to link stimulus responses to connectivity between brain areas.

Here we propose a framework that links stimulus responses to connectivity between brain areas. The framework is based on three quantities (1) External Stimulus Strength (ESS), which reflects the strength of the stimulus response in each brain area (2) External Stimulus Connectivity (ESC), which reflects connectivity between brain areas due to the stimulus responses and (3) Functional Connectivity (FC), which reflects connectivity both due to the stimulus response and due to internal (i.e. stimulus independent) activity. The three quantities allow us to address perceptual decoupling in a principled manner. Specifically, we hypothesized that during external cognition stimulus responses will be strong (high ESS) and connectivity between brain areas will largely reflect stimulus responses. In this case ESC will be similar (or coupled) to FC. During internal cognition however, stimulus responses will be attenuated (low ESS) and connectivity between brain areas will largely reflect internal processing. In this case, ESC will be decoupled from FC (Figure 1A).

**Figure 1.**
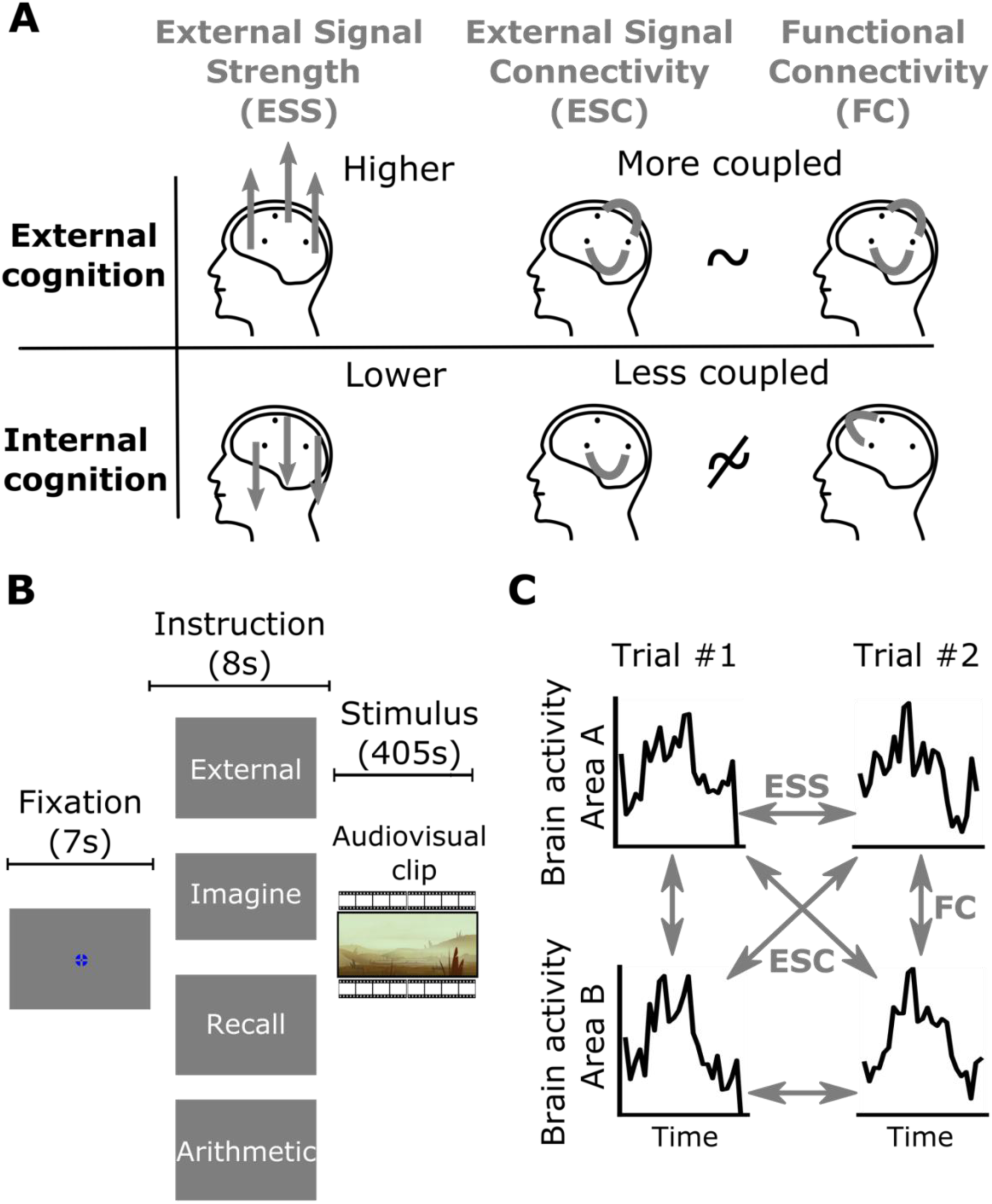
Conceptual framework and experimental paradigm. **A)** Framework for assessing the decoupling of brain processes during internal cognition. The framework is based on three quantities: External Signal Strength (ESS), External Signal Connectivity (ESC) and Functional Connectivity (FC). The figure depicts the three quantities (columns) during external and internal cognition (rows). The three dots inside each brain schematic represent hypothetical brain areas. ESS reflects the strength of the stimulus response in each brain area. ESC reflects the connectivity between brain areas due to the stimulus responses. FC reflects the connectivity between brain areas both due to the stimulus and due to internal (i.e. stimulus independent) activity. During external cognition the responses to external stimuli are strong (high ESS depicted as upward arrows) and connectivity between brain areas largely reflects stimulus responses. In this case, FC is coupled to ESC (the gray connectivity patterns are similar). During internal cognition, stimulus responses will be attenuated (low ESS depicted as downward arrows). In this case, connectivity between brain areas will generally reflect internal (stimulus independent) activity, such that FC will be decoupled from ESC (the gray connectivity patterns are different). **B**) Experimental paradigm. The figure represents a single trial (see Methods for details). We recorded brain activity using functional magnetic resonance imaging (fMRI) while presenting subjects with an audiovisual stimulus and instructing them to either engage in external or internal cognition. A trial began with a short fixation screen, followed by an instruction screen detailing whether to focus on the stimulus (external task) or engage in one of three internal tasks – imagery (imagine your next holiday), recall (recall your daily trip to work or school) or arithmetic (count backwards from 2000 in steps of 7). Note that the same audiovisual clip was presented for all tasks. Prior to the experiment, subjects were explicitly instructed to keep their eyes open and maintain fixation for all tasks. **C**) Analysis framework. We quantified ESS, ESC and FC as correlations within/across trials/areas. The figure represents area-averaged time courses in each of two hypothetical brain areas, A and B, for each of two trials. Arrows represent calculating the Pearson correlation coefficient between the time courses. We quantified ESS as the correlation between the time courses in each area across trials. This reflects the signal to noise ratio at each area. We quantified ESC as the correlation across areas and trials. This reflects the extent of shared stimulus processing across areas. We quantified FC in the usual way, as the correlation across areas in the same trial. Note that FC reflects a mixture of external stimulus responses and internal (i.e. stimulus independent) activity while ESS and ESC primarily reflect stimulus responses.

To test this, we measured whole brain activity using functional magnetic resonance imaging (fMRI) while subjects engaged in either external or internal cognition. Specifically, we presented subjects with a naturalistic audiovisual stimulus and instructed them to either engage in an external task (pay attention to the stimulus) or engage in one of three different internal tasks – mental arithmetic, recall or imagery. We chose these internal tasks based on previous studies (Ki et al., 2016; Shirer et al., 2012) and because their qualitatively different nature is likely to involve distinct functional connectivity patterns (Figure 1B, see Methods for details).

We analyzed the fMRI data by operationalizing the three quantities. We measured ESS as the correlation between the time courses in each brain area across trials. This quantifies the response reliability across repeated presentation of the stimulus and has previously been used as a measure of the signal to noise ratio (Hsu et al., 2004; Schoppe et al., 2016). We measured ESC as the correlation between the time courses across trials and areas. This generalization of the ESS quantifies the extent of shared stimulus processing across areas. Finally, we calculated FC in the standard way, that is as the correlation across areas in the same trial. Together, these definitions of ESC, ESS and FC provide a simple (mere correlations within/across trials/areas) and readily interpretable operationalization of our conceptual framework (Figure 1C, see Methods for details).

## Results

### Reduced External Signal Strength during internal cognition

Our conceptual framework suggests that during internal cognition stimulus responses will be attenuated and that this will be reflected in reduced ESS (Figure 1A). To test this, we recruited 20 subjects to complete the fMRI experiment. Each subject completed three trials for each of the four tasks, which allowed us to calculate ESS (as well as ESC and FC) for each task separately. Each trial began with a short fixation screen, followed by an instruction screen, followed by the presentation of the stimulus (Figure 1B). To analyze the data, we grouped voxels into a set of 61 previously defined ROIs. The 61 ROIs comprise 6 large-scale networks – Visual, Auditory/Language, DMN I, DMN II, Attention and Executive (Figure S1 and Methods) and were previously shown to reliably respond to narrative stimuli like our own (Regev et al., 2019). We quantified ESS as the Pearson correlation coefficient between the time courses in each brain area across trials (Figure 1C). For the external task, we expected reliable ESS across most ROIs. We found that all large-scale networks and 59 out of the 61 ROIs showed above chance ESS levels (assessed using group-level phase permutation tests, see Methods for details), indicating that much of the cortex is involved in stimulus processing for the external task (Figure 2A).

**Figure 2.**
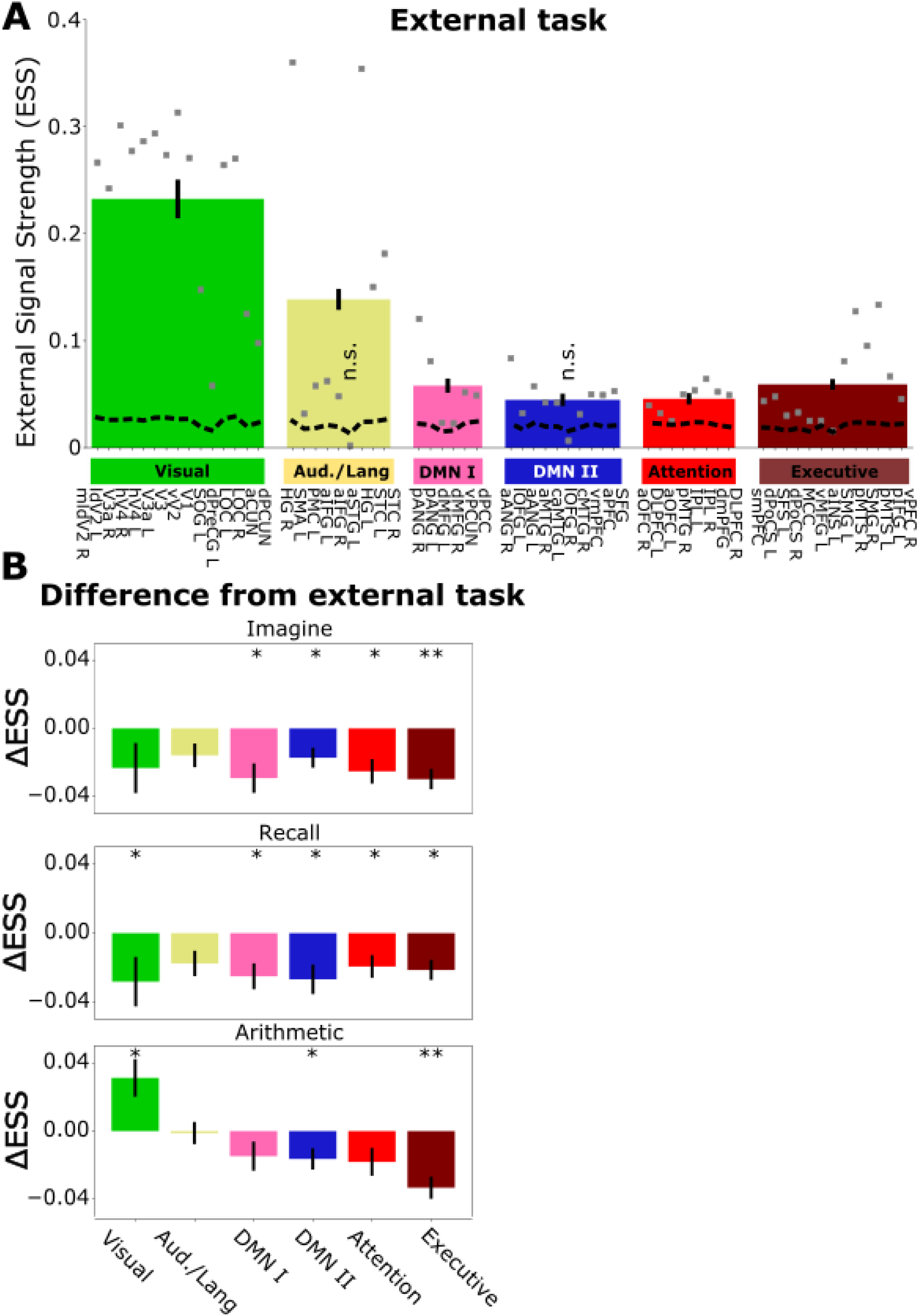
Reduced external signal strength (ESS) during internal cognition. **A**) ESS during external cognition. Each grey dot represents the ESS value at each of the 61 regions of interest (ROIs). The bars represent the average for each of the six large-scale networks. The black, dashed lines reflect the 97.5 percentile estimated using group-level phase permutation tests at each ROI (see Methods). The ESS is significantly above chance in all but two of the ROIs – left anterior superior temporal gyrus (aSTGL) and right lateral orbitofrontal gyrus (lOFG R) (see Methods and Figure S1 for details). **B**) Difference in ESS from the external task for each of the large-scale networks. The figure shows a general reduction except for the Visual network during arithmetic, which shows an increase. * and ** reflect significant reductions at the p<0.05 and p<0.01 level (Wilcoxon signed-rank tests with a false-discovery rate (FDR) adjustment across the 18 comparisons). Error bars represent SEM across subjects (N=20).

Next, we investigated how internal cognition affected these responses. We averaged the ESS across each of the large-scale networks and analyzed the difference from the external task. We found that the ESS was significantly lower for the DMN II and Executive networks for all three internal tasks. During imagine and recall, we further observed significant reductions in DMN I and the Attention networks. In the recall task we also observed a significant reduction in the Visual network. The Auditory/language network did not show a significant reduction in any of the tasks. Contrary to our expectations, we found a significant increase in ESS in the Visual network for the arithmetic task (Figure 2B).

One possibility is that the increase in the Visual network for the arithmetic task was due to the subjects failing to do the task and instead focusing on the stimulus. To rule this out we ran a separate experiment in which we behaviorally assessed the effects of the internal tasks on comprehension and recall of the stimulus. In this experiment subjects were allocated to four different groups, corresponding to the four tasks. Each subject viewed the stimulus while performing their allocated task. After completing the task, subjects were surprised with a computerized test assessing comprehension and recall of the stimulus. Test scores were significantly lower for all three internal tasks, confirming that subjects are concentrating on the internal tasks and not the stimulus (Figure S2). We return to the surprising increase in the Visual network for the arithmetic task in the Discussion.

### Functional connectivity patterns differentiate between tasks

The previous result demonstrates attenuated stimulus processing but does not reveal the brain processes that support each task. Based on previous work (Shirer et al., 2012) and in the context of our conceptual framework (Figure 1A), we expected that each task will involve distinct functional connectivity (FC) patterns. To test this, we computed the group-level FC matrices between the 61 ROIs for each task separately. These connectivity matrices suggested clear differences between the four tasks. For example, the imagine and recall tasks were marked by increased Visual-DMN II, DMN II-Attention and DMN II-Executive connectivity, whereas the arithmetic task was marked by reductions in these same connections (Figure 3A).

**Figure 3.**
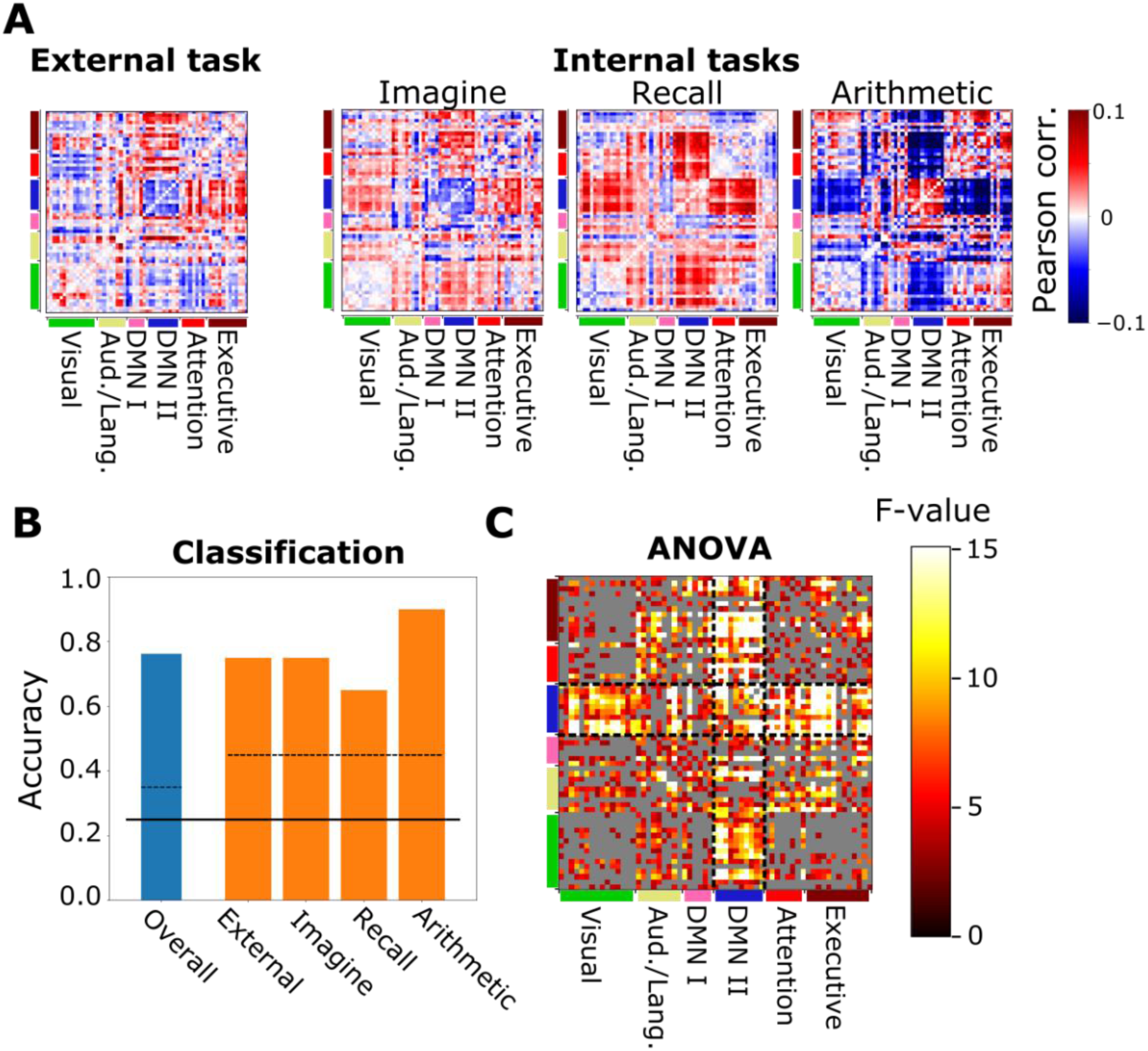
Functional connectivity robustly discriminates between the tasks. **A**) Group level FC matrices computed using all 61 ROIs for each of the tasks. The mean across all tasks was subtracted from each task in order to highlight the differences. **B**) Leave-one-subject-out classification results using the FC matrices as input. Dashed lines reflect 97.5 percentile of the null (binomial) distribution (see Methods). The high classification accuracy confirms that each task involves specific FC patterns. **C**) Repeated-measures ANOVA testing for the effect of task. The colors represent F-values for each connection. In total, 971 out of 1830 connections (53%) showed significant modulations (p<0.05 using FDR corrections, non-significant connections are shown in grey). 40 out of 45 of the connections (88.9%) within DMN II and 406 out of 510 connections (79.6%) between DMN II and the other networks were significantly modulated. Connections within DMN II and between DMN II and the other networks are enclosed by the dashed lines.

To assess the reliability of these differences we used leave-one-subject-out classification analysis. Each subject contributed four FC matrices as inputs, corresponding to each of the four tasks (see Methods for details). Overall mean classification accuracy was 0.76 (0.25 chance), indicating robust group level differences. Inspecting the classifier’s performance for each task separately confirmed that each task could be discriminated well above chance (0.75, 0.75, 0.65 and 0.9 for external, imagine, recall and arithmetic respectively). The confusion matrix did not suggest any systematic misclassifications, consistent with the high classification accuracy (Figure S3). Taken together, these results confirm that each task involves distinct FC patterns.

A limitation of the classification analysis is that although it provides a summary statistic of discriminability, it is opaque with respect to the involvement of each connection. To estimate how each connection is modulated we ran repeated measures ANOVA for each connection separately, testing for the effect of task. Based on previous work, we expected that connections between the DMN and Executive networks will be particularly strongly modulated (Andrews-Hanna et al., 2014; Benedek et al., 2016; Dixon et al., 2014; Kam et al., 2019; Smallwood et al., 2012; Turnbull et al., 2019; Zabelina & Andrews-Hanna, 2016). We found that most connections were significantly modulated by the tasks (53% of connections survived FDR correction at p<0.05). In particular, 88.9% (40/45) of connections within DMN II and 79.6% (406/510) of connections between DMN II and the other networks were significantly modulated, indicating that connectivity within and between this network is important for the tasks (Figure 3C).

### Functional connectivity is decoupled from the external stimulus during internal cognition

Our results so far show that internal cognition involves (1) attenuated stimulus responses (Figure 2B) and (2) specific FC patterns (Figure 3). However, the relationship between these two observations remains unclear. Based on our conceptual framework, we expected that during external cognition FC largely reflects stimulus processing. In this case, connectivity between brain areas solely due to the stimulus responses, termed External Signal Connectivity (ESC), will be similar (or coupled) to FC (Figure 1A). During internal cognition, FC will largely reflect internal (stimulus independent) processing. In this case, ESC will be different from FC, reflecting the decoupling of brain networks from stimulus processing.

To test this, we calculated the ESC matrices across all ROI pairs for each subject and task separately. Note that a useful property of the ESC matrix is that its diagonal corresponds to the ESS at each ROI, such that one may view the ESC as the generalization of ESS to between-area connectivity. To assess the similarity between the ESC and FC we calculated the Pearson correlation coefficient between the two matrices, which we refer to as FC-ESC coupling (Figure 4A, see Methods for details).

**Figure 4.**
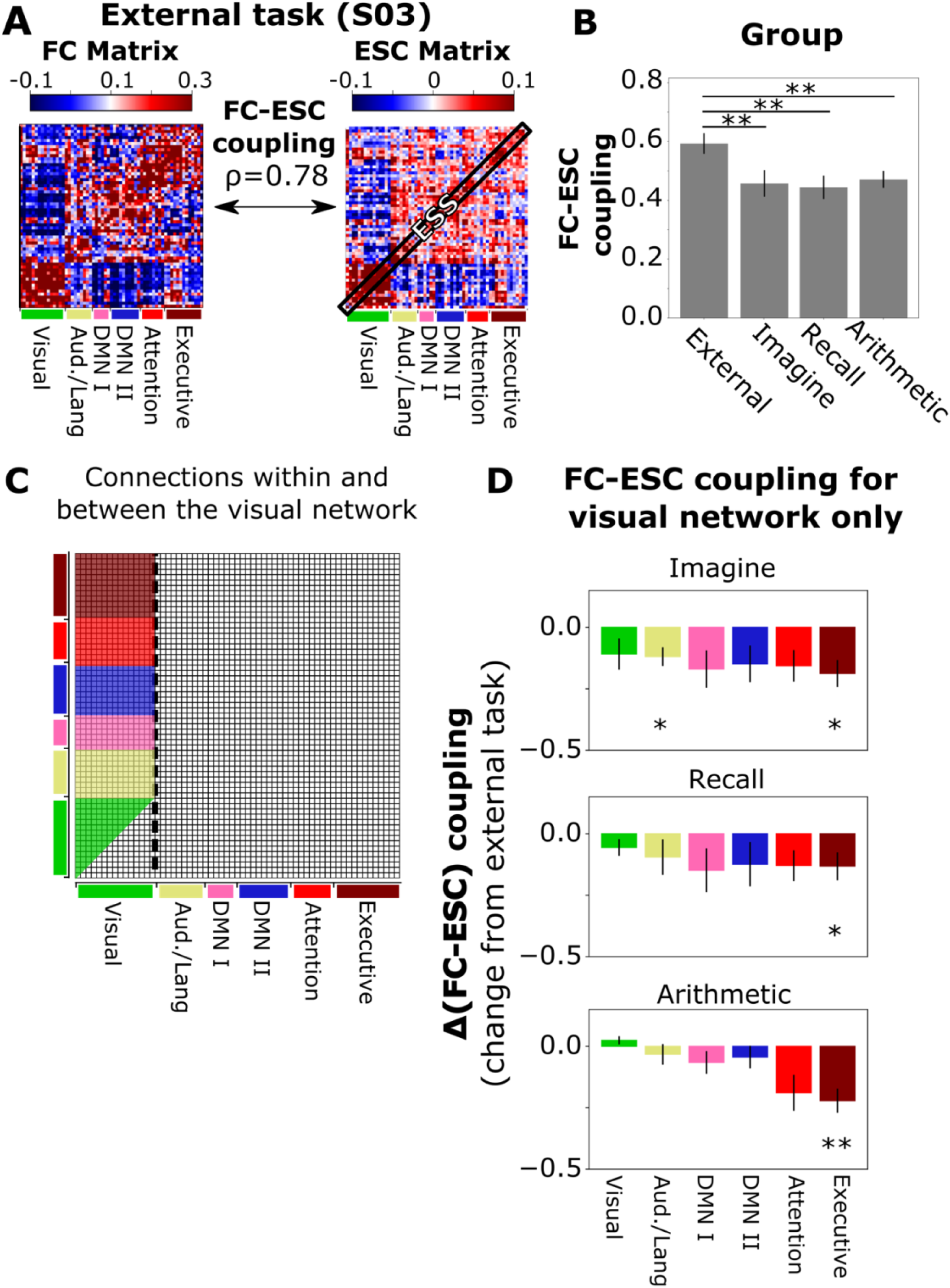
Reduced FC-ESC coupling during internal cognition. **A**) We assessed the coupling between FC and ESC by calculating the Pearson correlation coefficient between the FC and ESC matrices. The figure shows FC (left) and ESC (right) matrices for one subject (sub-03) for the external task. The high FC-ESC coupling (0.78) suggests that FC during external cognition primarily reflects the stimulus response. Note that the diagonal of the ESC matrix corresponds to the ESS at each ROI. **B**) Group-level FC-ESC is significantly higher for the external task as compared to each of the internal tasks. **C**) Connections used in calculating FC-ESC coupling within the Visual network and between the Visual network and the other networks (used in panel D). **D**) Change in FC-ESC coupling from the external task calculated using the connections in panel C. Visual-Executive FC-ESC coupling was lower for all internal tasks, even though the ESS in the Visual network increased for the arithmetic task (Figure 2B). Error bars represent SEM of the mean across subjects (N = 20). * and ** reflect p<0.05 and p<0.05 calculated using Wilcoxon signed-rank test and FDR adjustment.

Consistent with our hypothesis, we found that FC-ESC coupling was significantly higher for the external task as compared to each of the internal tasks (Figure 4B, Wilcoxon signed-rank tests for external task vs imagine, statistic = 26, p = 0.003, vs recall, statistic = 34, p=0.008, vs arithmetic, statistic = 28, p=0.004). This quantitively demonstrates that brain networks are decoupled from the stimulus during internal cognition.

Finally, we reexamined the unexpected increase in ESS in the Visual network for the arithmetic task (Figure 2B). Specifically, we speculated that even though the strength of the response increased (ESS increased) in the Visual network, FC-ESC coupling actually reduced. To test this, we re-calculated FC-ESC coupling separately for each large-scale network and using only connections with the Visual network (Figure 4C). We found reduced FC-ESC coupling between the Visual and Executive networks for all three internal tasks (Figure 4D). Thus, even though stimulus responses in the Visual network increased for the arithmetic task, connectivity between the Executive and Visual networks became more reflective of internal (i.e. stimulus independent) activity than stimulus processing.

## Discussion

Here we proposed a conceptual and analysis framework to assess the decoupling of brain processes from the external stimulus during internal cognition (Figure 1A). We tested this framework using an fMRI experiment in which subjects engaged either in external or internal tasks (Figure 1B). We found that for the internal tasks stimulus responses were generally reduced (Figure 2), that each task was characterized by specific FC connectivity patterns (Figure 3), and that these FC patterns were less coupled to the stimulus as compared to the external task (Figure 4). Together, our results quantitively demonstrate that brain processes are decoupled from the external stimulus during internal cognition.

### Conceptual and analysis framework

A major contribution of this paper is the conceptual and analysis framework for assessing decoupling from the external stimulus (Figure 1A, Figure 1C). The key feature of the framework is that it links stimulus processing to connectivity between brain areas, thus providing a quantitative measure of decoupling from the external stimulus. We carefully constructed the framework to be both simple and interpretable, only requiring the computation of correlations within/across trials/areas. From an experimental viewpoint, the only requirement for using this framework is that the stimulus is presented more than once, which is standard practice in neuroscience. This means that the framework can be easily adopted by future studies. While other ways of linking stimulus responses to connectivity between brain areas are possible (e.g. by explicitly modelling the response to the stimulus), these approaches are more complicated, difficult to interpret and potentially require tailored experimental paradigms.

### Reduced External Signal Strength

Our finding of reduced ESS during internal cognition is consistent with previous EEG studies that showed reduced stimulus responses during mind wondering (Baird et al., 2014; Barron et al., 2011; Kam et al., 2011; Schooler et al., 2011). Our study goes further by investigating these reductions across different brain networks. This showed that while stimulus responses are generally reduced, they can increase in some cases (as was the case for the Visual network for the arithmetic task, see below). Thus, a reduction in stimulus responses may not be a universal feature of internal cognition. Notwithstanding this unexpected finding, we found significant reductions in Executive and DMN II networks for all the internal tasks (Figure 2B). While it is tempting to interpret this as evidence for the importance of these networks, especially in light of the strong modulation of DMN II connectivity by the tasks (Figure 3C), we did not carry out any across-area comparisons. The reason for this is that each ROI reflects a different number of voxels, and there are different numbers of ROIs in each network. Proper comparisons between areas will need to carefully account for these differences, which is outside the scope of this paper.

### Unexpected increase in External Signal Strength for the Visual network during arithmetic

We unexpectedly found increased ESS in the Visual network for the arithmetic task. This finding appears generally incompatible with the notion of perceptual decoupling and is at odds with a previous EEG study that used this task and found reduced stimulus processing (Ki et al., 2016). In a separate behavioral study, we ruled out that this is due to the subjects not performing the task (Figure S2). With respect to the Ki et al study, the source of the discrepancy is unclear due differences in recording (here fMRI vs EEG in Ki et al) and analysis method (here ESS vs a measure of reproducibility across subjects in Ki et al), but one potential explanation is that the brain-wide approach taken in that study combined with the coarse spatial resolution of EEG meant that the increase in the Visual network was lost amongst the reductions in other areas.

### Decoupling from the external stimulus during internal cognition

We showed that (1) each task involves specific FC patterns (Figure 3) and (2) that these FC patterns are less reflective of the stimulus during internal cognition (Figure 4). This provides a quantitative demonstration that brain processes are decoupled from external stimuli during internal cognition. This finding was made possible by our framework which quantitively links stimulus processing to connectivity between brain areas. This quantitative approach also allowed us to reconsider the arithmetic task, for which stimulus response increased in the Visual network (Figure 2B). Specifically, in spite of the stronger responses in the Visual system, connectivity between the Visual and Executive networks was less coupled to the stimulus. Decoupling with the Executive network may be particularly important in our experiment since our internal tasks are “top-down” (sometime referred to as cued, intentional or volitional (Dixon et al., 2014)) and are thus likely to involve executive processes. It would be interesting to examine if a similar decoupling of connectivity is observed during spontaneous forms of internal cognition such as mind-wandering and daydreaming.

## Methods

### Subjects

We recruited 24 subjects for the fMRI study. Two subjects opted out before completing the experiment and two were rejected from analysis due to excessive movement (median framewise displacement > 0.2 (Vanderwal et al., 2017)). Thus, we analyzed data from a total of 20 subjects (9F, ages 20-37, median 27). For the behavioral study (Figure S2) we recruited a separate group of 32 subjects (10F, ages 20-47, median 22).

We obtained informed consent from all subjects. The experimental protocol was approved by the ethics and safety committees of the National Institute of Information and Technology, Japan.

### Experimental design

Subjects viewed an audiovisual stimulus while engaging in either an external task or one of three internal tasks (Figure 1A). For the external task, subjects were instructed to pay careful attention to the audiovisual stimulus. In the internal tasks, subjects were instructed to try to ignore the stimulus and either imagine their next holiday (imagine), recall their daily trip to work or school (recall) or count backwards from 2000 in steps of 7 (arithmetic). For the imagine task, subjects were explicitly instructed to imagine a holiday that has not yet occurred, otherwise subjects may recall a previous holiday. For the internal tasks, subjects were also instructed to (1) perform all tasks in their head (as opposed to out loud), (2) recall and imagine in as much detail as possible and (3) continue as best they can in case of a mistake or lapse in attention. For all tasks, subjects were also instructed to keep their eyes open and fixated.

We chose the arithmetic and recall tasks because they have been used in other studies (Ki et al., 2016; Shirer et al., 2012). We added the imagine task as a future-oriented, divergent form of internal cognition. Together, the three tasks reflect qualitatively different forms of internal cognition.

### Audiovisual stimulus

For the stimulus we used naturalistic audiovisual narratives, which are known to elicit reliable, brain-wide responses (Hasson et al., 2010). The stimulus consisted of 7 short audiovisual clips (each clip lasted 30 - 225s) of animations and natural movies lasting a total of 13.5minutes (one subject, S02, viewed a longer version of the stimuli in which the final clip was two minutes longer, resulting in 15.5minutes in total. We truncated this subject’s data to 13.5 minutes in order to conform with the rest of the subjects). The videos were sourced from Vimeo (The Black Hole, A Single Life, How They Get There, Gregory Is a Dancer, Grounded, Trashonauts). The clips were cropped and grouped into two different sequences lasting 405s each. We opted for clips with narrative since these also involve the activation of higher order areas such as the DMN (Simony et al., 2016), and relatively little linguistic content in order to minimize the effect of the bilingual nature of the subjects (international students studying in Japan).

### fMRI experiment

Each of the 20 subjects completed the experiment across three sessions. During each session, subjects viewed each of the two audiovisual sequences while performing each of the four tasks, resulting in eight trials in total (recorded in separate runs) per subject. Thus, across the three sessions, each subject completed 24 trials (4 tasks x 2 stimulus sequences x 3 repeats). Each trial began with the presentation of a fixation screen (7 seconds), followed by the instructions for that trial (8 seconds, i.e. whether to externally attend, imagine, recall or count backwords), followed by the presentation of the stimulus (Figure 1A). The task order was randomized with the restrictions that (1) tasks repeated twice during each session (2) tasks did not repeat consecutively. The two stimulus sequences alternated consecutively. A high contrast anatomical image was acquired in one of the three sessions.

The stimulus was presented through a projector screen inside the scanner at a resolution of 1920×1080 at 60Hz (visual angle ~22.0×12.5) using the Presentation software (Neurobehavioral Systems, Albany, CA, USA). A fixation cross (35×35 pixels) was presented throughout. Sound was relayed using MR-compatible earphones.

Functional scans were collected on a 3T Siemens Prisma scanner (Siemens, Germany) using a 64-channel coil and a multiband gradient echo-planar imaging (EPI) sequence (Moeller et al., 2010; repetition time (TR) = 1000 ms; echo time (TE) =30 ms; flip angle = 62°; voxel size = 2×2×2 mm; matrix size =96×96; field of view (FOV) = 192 x 192 mm; multiband factor = 3). Seventy-two axial slices covered the entire cortex. Anatomical data were collected using a T1-weighted magnetization-prepared rapid acquisition with gradient echo (MPRAGE) sequence (TR = 2530 ms; TE = 3.26 ms; flip angle = 9; voxel size = 1×1×1 mm; matrix size = 256×256; FOV=256×256 mm) on the same 3T scanner.

### Behavioral experiment

We ran a separate behavioral experiment to validate that the subjects were performing the internal tasks and not focusing on the stimulus. We recruited a separate group of 32 subjects (see Subjects) and divided them into four groups of 8, corresponding to the four tasks. Each subject viewed the audiovisual stimulus while performing the task assigned to their group. Subjects were then surprised with a computerized test assessing comprehension and recall of the stimulus. If the subjects were correctly performing the tasks and concentrating internally, then test scores should be lower for the internal task groups vs the external task group.

The instructions and stimulus were identical to the fMRI task, but the presentation order was slightly different. Each experiment began with an instruction screen, (i.e. whether to externally attend, imagine, recall or count backwards; same as in Figure 1A), followed by a 15s practice stimulus (from a separate clip). Subjects had the opportunity to seek clarification before continuing to the first stimulus sequence (405s). After an optional short break, the subjects viewed the second sequence (405s). After viewing both sequences subjects were surprised with a computerized, 30 question, 4-option multiple choice test. The questions varied from specific detail (e.g. “In the clip where the man gets hit by the car, what does the man drink in the start of the clip?”) to narrative related question (“In the clip where the man gets hit by the car, why does he get hit by a car?”). Subjects had 14.5s to answer each question. A warning was displayed with 5s remaining. The stimulus and multiple choice test were presented using PsychoPy 3.2.4 (Peirce et al., 2019) on a 15inch Macbook pro (4^th^ generation, ~32.2×22.4° of visual angle).

### fMRI preprocessing

All MRI data in this manuscript were preprocessed using fmriprep. Below is the fmriprep-generated boilerplate.

Start of boiler plate:

Results included in this manuscript come from preprocessing performed using fMRIPrep 1.4.1 (Esteban, Blair, et al., 2018; Esteban, Markiewicz, et al., 2018), which is based on Nipype 1.2.0 (K. Gorgolewski et al., 2011; K. J. Gorgolewski et al., 2018).

#### Anatomical data preprocessing

The T1-weighted (T1w) image was corrected for intensity non-uniformity (INU) with N4BiasFieldCorrection (Tustison et al., 2010), distributed with ANTs 2.2.0 (Tustison et al., 2010) and used as T1w-reference throughout the workflow. The T1w-reference was then skull-stripped with a Nipype implementation of the antsBrainExtraction.sh workflow (from ANTs), using OASIS30ANTs as target template. Brain tissue segmentation of cerebrospinal fluid (CSF), white-matter (WM) and gray-matter (GM) was performed on the brain-extracted T1w using fast (FSL 5.0.9, RRID:SCR_002823, (Zhang et al., 2001)). Brain surfaces were reconstructed using recon-all (FreeSurfer 6.0.1, (Dale et al., 1999)), and the brain mask estimated previously was refined with a custom variation of the method to reconcile ANTs-derived and FreeSurfer-derived segmentations of the cortical gray-matter of Mindboggle (RRID:SCR_002438, (Klein et al., 2017)). Volume-based spatial normalization to one standard space (MNI152NLin2009cAsym) was performed through nonlinear registration with antsRegistration (ANTs 2.2.0), using brain-extracted versions of both T1w reference and the T1w template. The following template was selected for spatial normalization: ICBM 152 Nonlinear Asymmetrical template version 2009c [(Fonov et al., 2009), RRID:SCR_008796; TemplateFlow ID: MNI152NLin2009cAsym].

#### Functional data preprocessing

For each of the 24 BOLD runs found per subject (across all tasks and sessions), the following preprocessing was performed. First, a reference volume and its skull-stripped version were generated using a custom methodology of fMRIPrep. The BOLD reference was then co-registered to the T1w reference using bbregister (FreeSurfer) which implements boundary-based registration (Greve & Fischl, 2009). Co-registration was configured with nine degrees of freedom to account for distortions remaining in the BOLD reference. Head-motion parameters with respect to the BOLD reference (transformation matrices, and six corresponding rotation and translation parameters) are estimated before any spatiotemporal filtering using mcflirt (FSL 5.0.9, (Jenkinson et al., 2002)). BOLD runs were slice-time corrected using 3dTshift from AFNI 20160207 ((Cox & Hyde, 1997), RRID:SCR_005927). The BOLD time-series, were resampled to surfaces on the following spaces: fsaverage5. The BOLD time-series (including slice-timing correction when applied) were resampled onto their original, native space by applying a single, composite transform to correct for head-motion and susceptibility distortions. These resampled BOLD time-series will be referred to as preprocessed BOLD in original space, or just preprocessed BOLD. The BOLD time-series were resampled into standard space, generating a preprocessed BOLD run in [‘MNI152NLin2009cAsym’] space. First, a reference volume and its skull-stripped version were generated using a custom methodology of fMRIPrep. Several confounding time-series were calculated based on the preprocessed BOLD: framewise displacement (FD), DVARS and three region-wise global signals. FD and DVARS are calculated for each functional run, both using their implementations in Nipype (following the definitions by (Power et al., 2014)). The three global signals are extracted within the CSF, the WM, and the whole-brain masks. Additionally, a set of physiological regressors were extracted to allow for component-based noise correction (CompCor, (Behzadi et al., 2007)). Principal components are estimated after high-pass filtering the preprocessed BOLD time-series (using a discrete cosine filter with 128s cut-off) for the two CompCor variants: temporal (tCompCor) and anatomical (aCompCor). tCompCor components are then calculated from the top 5% variable voxels within a mask covering the subcortical regions. This subcortical mask is obtained by heavily eroding the brain mask, which ensures it does not include cortical GM regions. For aCompCor, components are calculated within the intersection of the aforementioned mask and the union of CSF and WM masks calculated in T1w space, after their projection to the native space of each functional run (using the inverse BOLD-to-T1w transformation). Components are also calculated separately within the WM and CSF masks. For each CompCor decomposition, the k components with the largest singular values are retained, such that the retained components’ time series are sufficient to explain 50 percent of variance across the nuisance mask (CSF, WM, combined, or temporal). The remaining components are dropped from consideration. The head-motion estimates calculated in the correction step were also placed within the corresponding confounds file. The confound time series derived from head motion estimates and global signals were expanded with the inclusion of temporal derivatives and quadratic terms for each (Satterthwaite et al., 2013). Frames that exceeded a threshold of 0.5 mm FD or 1.5 standardised DVARS were annotated as motion outliers. All resamplings can be performed with a single interpolation step by composing all the pertinent transformations (i.e. head-motion transform matrices, susceptibility distortion correction when available, and co-registrations to anatomical and output spaces). Gridded (volumetric) resamplings were performed using antsApplyTransforms (ANTs), configured with Lanczos interpolation to minimize the smoothing effects of other kernels (Lanczos, 1964). Non-gridded (surface) resamplings were performed using mri_vol2surf (FreeSurfer).

Many internal operations of fMRIPrep use Nilearn 0.5.2 ((Abraham et al., 2014), RRID:SCR_001362), mostly within the functional processing workflow. For more details of the pipeline, see the section corresponding to workflows in fMRIPrep’s documentation.

End of boiler plate.

### Confound removals

We used the fmriprep-estimated confounds to regress out head movement, their temporal derivatives, and their squares (24 DOF in total). We further included the top 5 components from PCA decompositions of the white matter and cerebrospinal fluid voxels. This is the “24HMP + aCompCor” denoising pipeline of (Parkes et al., 2018). We also high-passed the data above 0.01Hz and removed the first 40s of each run from the analysis. Confound removal was carried out by a single call to nilearn’s signal.clean() routine (Abraham et al., 2014).

### Regions of interest

We analyzed the data using the 61 ROIs defined in (Regev et al., 2019). Regev et al investigated the effects of visual vs auditory attention on narrative stimulus processing and reported clear differences in most of these ROIs. We reasoned that these ROIs will also respond to our narrative stimuli. The 61 ROIs comprise six large-scale networks: Visual, Auditory/Language, DMN I, DMN II, Attention and Executive. Note that DMN I and DMN II are not based on the “core” and related subdivisions of the DMN as defined in (Andrews-Hanna et al., 2010, 2014). We used pycortex (Gao et al., 2015) to display the ROIs on top of a flattened cortex (fsaverage, Figure 1S)

All analysis in this manuscript are based on the 61 ROIs time courses, obtained by spatial averaging across all voxels in each ROI. Further information about the ROIs can be found in the original publication (Regev et al., 2019).

### External Signal Strength, External Signal Connectivity and Functional Connectivity

Our analysis framework involves three key quantities: External Signal Strength (ESS), External Signal Connectivity (ESC) and Functional Connectivity (FC). We calculated ESS as the Pearson correlation coefficient between the time courses in the same ROI across repeated presentations of the stimulus. This definition of the ESS, sometime referred to as signal to noise ratio or noise ceiling, is used for estimating the maximum performance of encoding models (e.g. (Huth et al., 2012, 2016)). We calculated FC in the usual way, as the correlation between the time courses across areas in the same trial (Figure 1C).

To relate the ESS to FC we were inspired by recent developments in inter-subject correlation analysis, where the activity in a given brain area from one subject is correlated with the activity in another brain area from a different subject. This reflects the extent of shared stimulus processing in the two areas (Regev et al., 2019; Simony et al., 2016). We adopted this approach to our paradigm by calculating the correlation between the time course in one ROI and a given trial and the time course in another ROI in a different trial (Figure 1C). Because this procedure is not symmetric (correlation between the time courses in area A for trial N and area B for trial M is generally different from the correlation between the time courses in area B for trial N and area A for trial M), we averaged across both estimates (similar to (Regev et al., 2019; Simony et al., 2016)). We refer to this quantity as External Stimulus Connectivity (ESC). An ESC matrix is calculated by repeating the computation across all ROI pairs. The diagonal of this matrix corresponds to the ESS in each area, consistent with viewing ESC as a generalization of the ESS computation (Figure 4A).

In our experimental paradigm each subject viewed the stimulus three times for each of the four tasks. To calculate FC for each task we concatenated the time courses across the three repeats and calculated the correlation across all ROI pairs. Specifically, denoting the time course for ROI *i* ∈ {1… 61} at trial *j* ∈ {1… 3} as 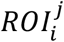, we concatenated the time courses across the three repeats, denoted 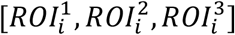, and calculated the Pearson correlation coefficient across all area pairs 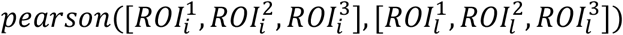, *i,l* ∈ {1…61}. To calculate ESC, we simply shifted the concatenated time course in one of the ROIs by one trial and repeated the procedure 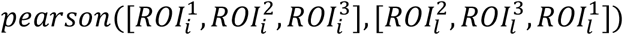, *i,l* ∈ {1…61} (note the superscripts in the second term). We note that the best way of estimating the ESS for finite data is the subject of ongoing research and our definition differs from that of others (Hsu et al., 2004; Lage-Castellanos et al., 2019; Schoppe et al., 2016). The advantage of our concatenation + shift approach is that it only involves a minimal change to the way in which we calculate FC, making comparison between these two quantities more natural.

ESS, ESC and FC are calculated using the same data and are thus related in some way. One concern is that changes in one quantity (e.g. reductions in ESS) are trivially predictive of changes in the other quantities (e.g. reductions in ESC or FC). However, from an analytic perspective it is straightforward to simulate systems in which changes in one quantity do not have a simple relationship to the changes in the other quantities (Figure S4). From an empirical perspective, we observed that both a reduction and increases in ESS (Figure 2B, imagine and recall vs arithmetic) resulted in a decrease of FC-ESC coupling (Figure 4D). Thus, our FC-ESC decoupling results are not trivially determined by the changes in ESS.

### Phase randomization

We used phase randomization to estimate the group-level significance of the ESS estimates in the external task (Figure 2A). We created a surrogate dataset by randomizing the phase of each ROI time course for each subject and trial. We then repeated the group-level ESS calculation using this surrogate data set. This procedure was repeated 2000 times to construct a null distribution which we used to obtain p-values. We corrected for multiple comparisons across ROIs using the false-discovery rate with alpha=0.05 (Yekutieli & Benjamini, 1999). Two of the 61 ROIs did not survive this procedure (Figure 2A).

### Classification analysis

We used classification analysis to assess whether FC patterns can reliably distinguish between the four tasks (Figure 3). We used a-leave-one-subject-out procedure where the data from all subjects except one is used to train a classifier and the data from the held-out subject is used to test the classifier. The input to the classifier was a 1830 dimensional vector composed of the upper triangle of the FC matrices. Each subject contributed four such vectors, corresponding to the four tasks. As a classifier we used a linear support vector machine with the default parameters (C=1) in the scikit-learn (v0.22.1) implementation (Pedregosa et al., 2011).

We report classification accuracy as the average across all held-out subjects (N=20). To assess significance, we calculated confidence intervals using the binomial distribution. For the overall accuracy (purple bar in Figure 3B) we used probability of success = 0.25 (four tasks) and n=80 (20 subjects x 4 tasks). The 98.3 percentile corresponds to 28/80=0.35. When separating the accuracy for each task (orange bars in Figure 3B) we used the binomial distribution with probability of success = 0.25 and n = 20, for which the 98.6 interval corresponds to 9/20=0.45.

### Statistics

For statistics we used a combination of non-parametric tests, phase randomization and ANOVA as described above and in the main text. When assessing the difference in ESS (Figure 2B, Wilcoxon sign-rank tests) and when conducting the repeated measures ANOVA (Figure 3C) we Fisher z-transformed the raw correlations values before conducting the tests. We also applied the Fisher z-transform before calculating the FC-ESC coupling (Cole et al., 2014).

We used scipy (v1.3.2) to run the non-parametric tests (Mann-Whitney and Wilcoxon signed-rank). For the FDR adjustment and ANOVA we used stats-models (v0.11.0). We used custom python code to implement the phase randomization.

## Online availability

Preprocessed data and code to run all analysis are available online at DOI 10.17605/OSF.IO/47×EB.

## Acknowledgments

We thank Janice Chen for stimulus suggestions. DC was supported by a JSPS international postdoctoral research fellowship, JSPS KAKENHI18F18706, and JST CREST JPMJCR18A5. SN was supported by JST CREST JPMJCR18A5 and ERATO JPMJER1801. TN was supported by JSPS KAKENHI Grant Number JP20K07718 and JP20H05023 in #4903 (Evolinguistics).

## Supplementary material

**Figure S1.**
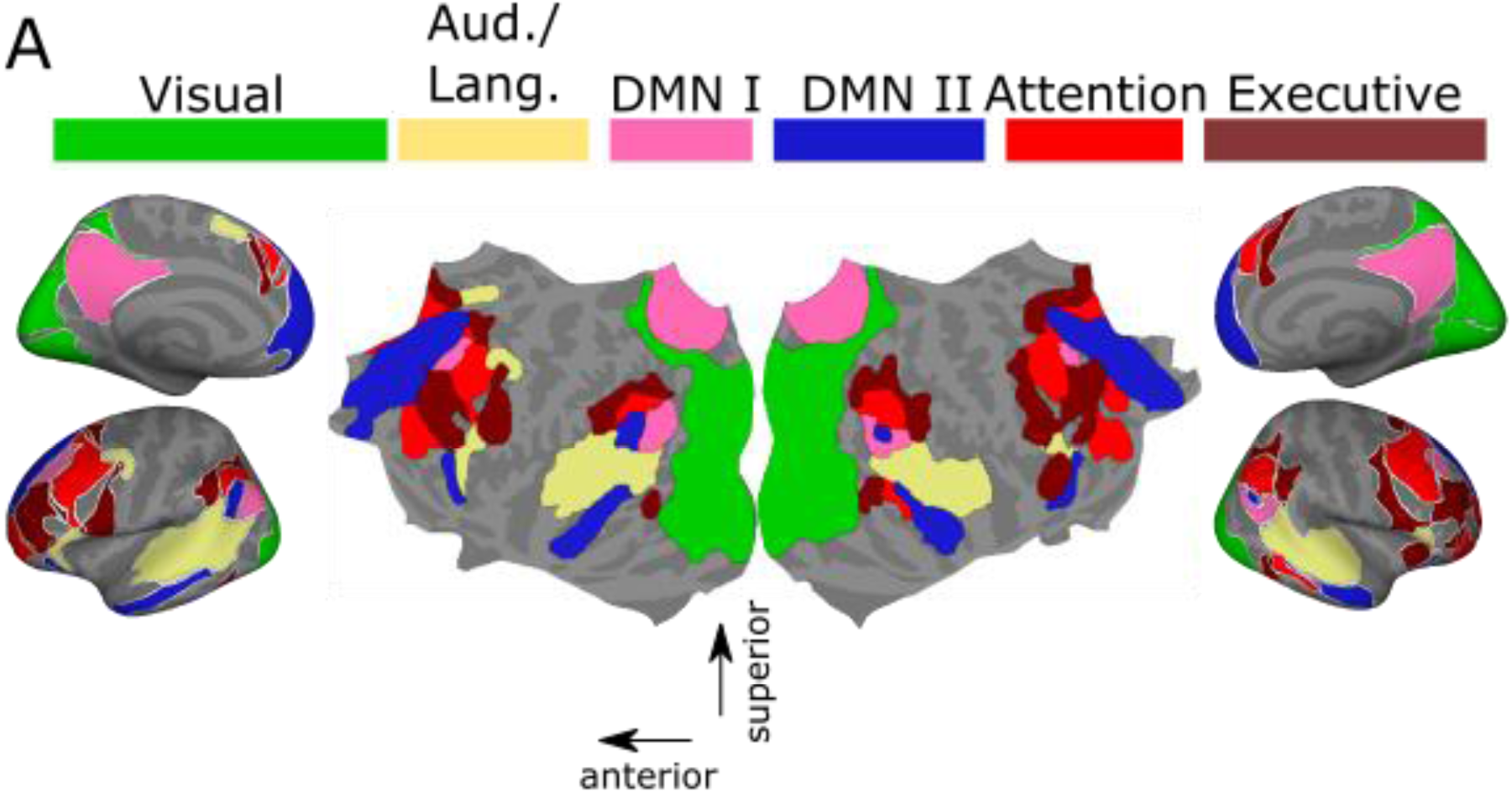
ROIs used in the analysis. The figure shows the 6 large-scale networks on a flattened cortex along with inflated medial and lateral views. See (Regev et al., 2019) for detailed description of the ROIs and their coordinates.

**Figure S2.**
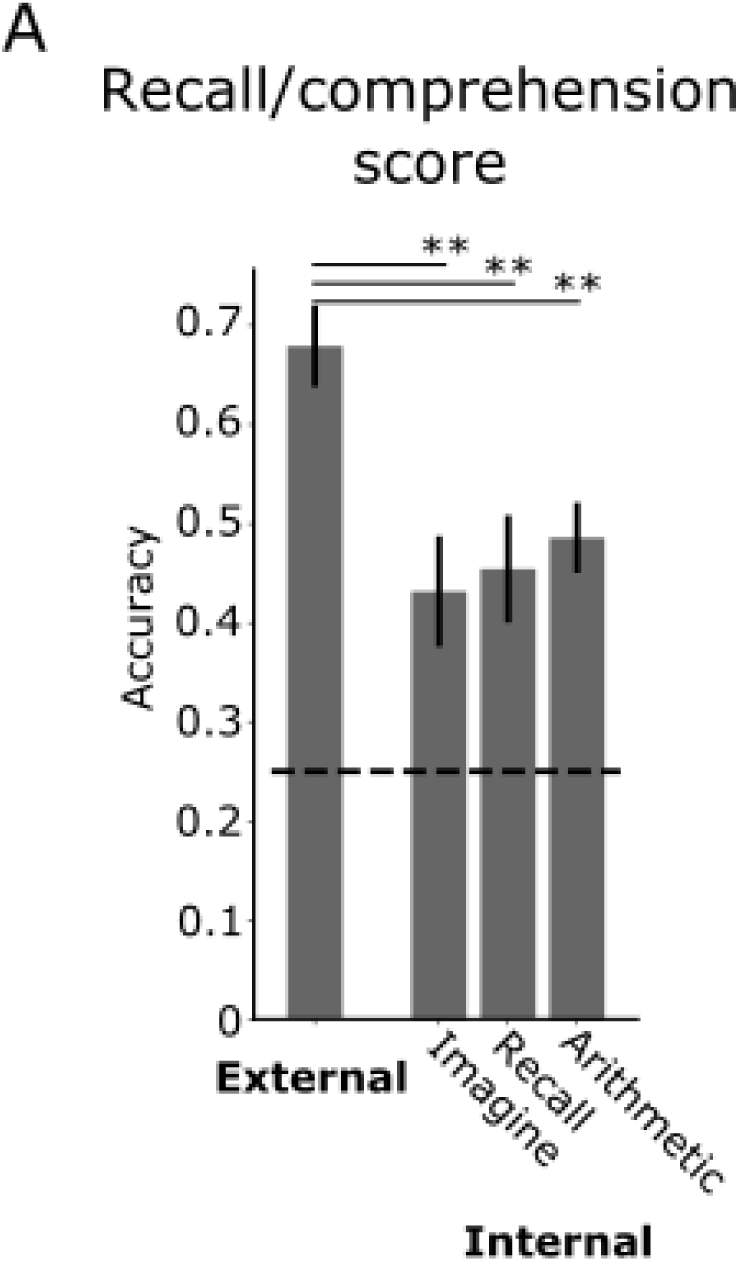
Reduced stimulus comprehension and recall for the internal tasks. 32 subjects were assigned to four groups (8 in each group) corresponding to the four tasks. Each subject viewed the stimulus while performing the task allocated to their group. Subjects were then surprised with a multiple choices test (4-choice, 30 questions in total) assessing comprehension and recall of the stimulus. The group performing the external task scored significantly higher on the test as compared to each of the internal task groups (Mann-Whitney tests for external vs imagine, U = 8, p = 0.006, vs recall, U = 5.5,p=0.0029, vs arithmetic, U = 5, p=0.002). Error bars reflects SEM (N=8).

**Figure S3.**
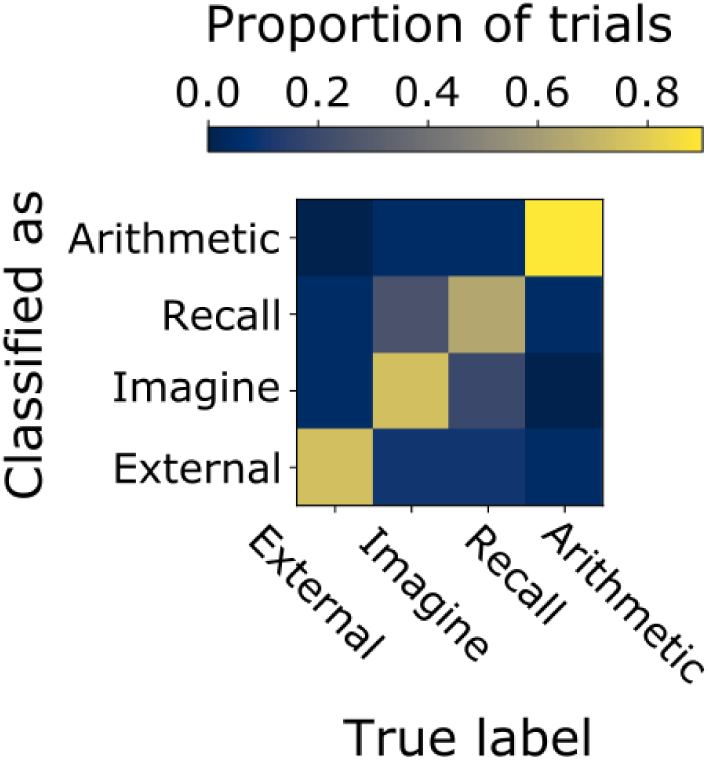
Confusion matrix for the classification analysis.

**Figure S4.**
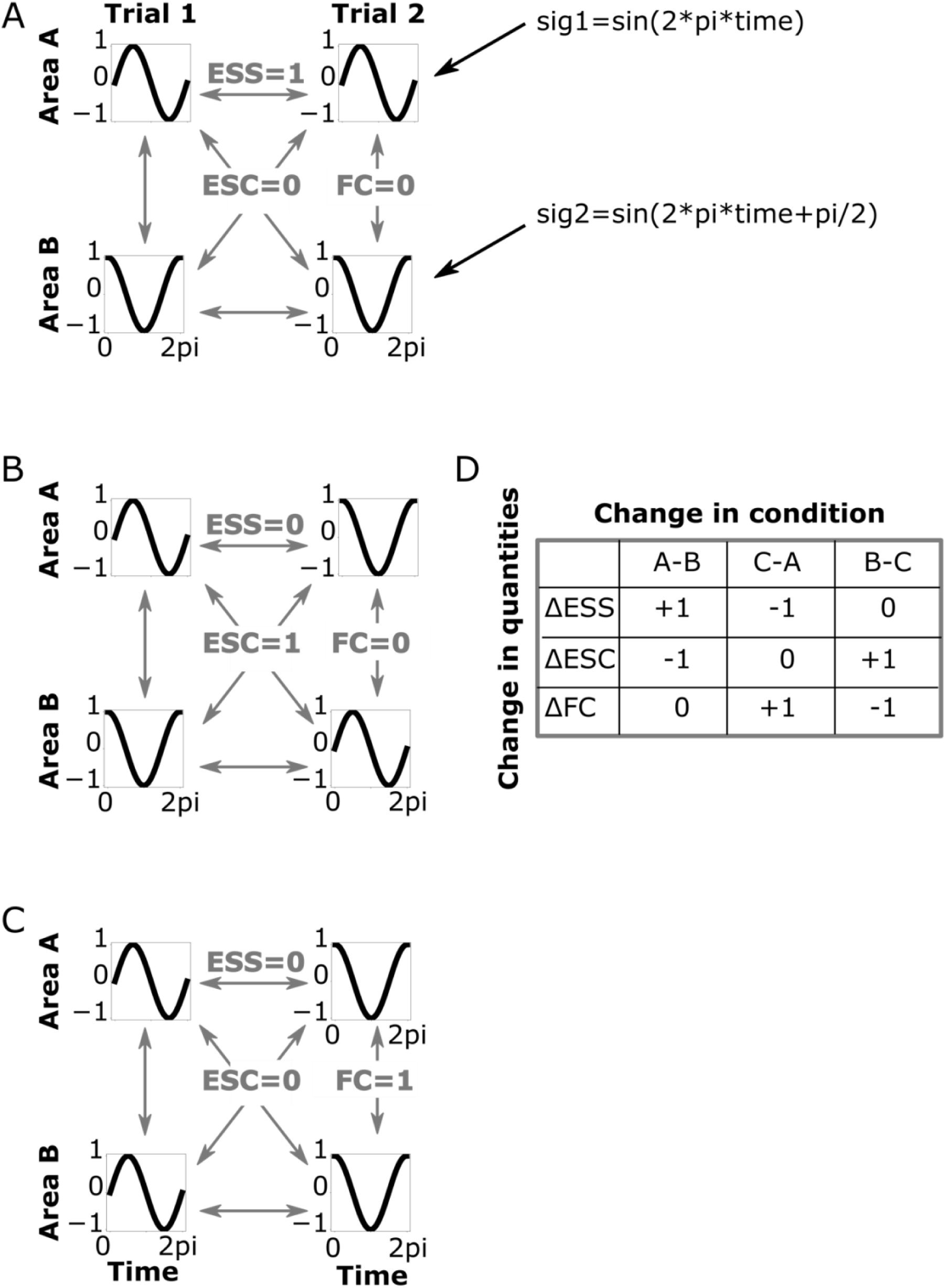
A change in one of the quantities (ESS, ESC and FC) is not trivially predictive of changes in the other quantities. We generated two sinusoidal signals, sig1 = sin(2*pi*time) and sig2 = sin(2*pi*time+pi/2), such that corr(sig1,sig2) = 0, and assigned them differently across areas and trials, resulting in different values for ESS, ESC and FC. **A**) The time coursein area A for both trials is sig1 and the time course in area B for both trials is sig2. ESS = 1, ESC =0 and FC = 0. **B**) The time course in area A for trial 1 and area B for trial 2 is sig1. The time course in area B for trial 1 and area A for trial 2 is sig2. ESS = 0, ESC = 1 and FC =0. **C**) The time course in areas A and B for trial 1 is sig1. The time course in areas A and B for trial 2 is sig2. ESS = 0, ESC = 0 and FC = 1. **D**) Changes in the quantities between the scenarios. Even in this very simple scenario (only two signals as opposed to four and no noise), a change in one quantity is not trivially predictive of changes in the other quantities.

